# The ATLAS™ screening assay reveals distinct CD4^+^ and CD8^+^ SARS-CoV-2 antigen response profiles which have implications to Omicron cellular immunity

**DOI:** 10.1101/2022.05.17.491668

**Authors:** James J. Foti, Kevin Lema, Justin Strickland, Emily Tjon, Adrienne Li, Amalia Rivera, Crystal Cabral, Laura Cormier, Louisa Dowal, Sudhir Rao, Vijetha Vemulapalli, Jessica B. Flechtner

## Abstract

The emergence of SARS-CoV-2 variants are a persistent threat to the efficacy of currently developed prophylactic vaccines and therapeutic antibodies. These variants accumulate mutations in the spike protein which encodes the epitopes necessary for neutralizing antibody binding. Moreover, emerging evidence suggest that robust antibody responses are insufficient to prevent severe disease and long-lasting viral immunity requires T cells. Thus, understanding how the T cell antigen landscape evolves in the context of these emerging variants remains crucial. T cells responses are durable and recognize a wider breadth of epitopes reducing the possibility of immune escape through mutation. Here, we deploy the ATLAS™ assay which identifies CD4^+^ and CD8^+^ T cell antigens by utilizing the endogenous HLA class-I and class-II peptide processing pathways. Profiling of T cells from exposed and unexposed donors revealed rich and complex patterns which highlighted the breadth of antigenic potential encoded in SARS-CoV-2. ATLAS revealed several common or frequent antigenic regions as well as an abundance of responses in the unexposed cohort potentially the result of pre-exposure to related coronaviruses. ORF10 was a common CD4^+^ response in the unexposed cohort while spike was identified as a common and frequent target in both cohorts. Moreover, the spike response profiles allowed us to accurately predict the impact of Omicron spike mutations. This analysis could thus be applied to study the impact of future emerging VOCs.

## Introduction

At the time of writing this report, just over two years have passed since the beginning of the COVID-19 pandemic. Globally, SARS-CoV-2, the etiologic agent of the disease, has taken a human toll approaching the 5.5 million mark, with numbers continuing to rise due to periodic infection waves linked to seasonal cycles and the emergence of fast-spreading viral Variants of Concern (VOC). VOCs are viral strains that accumulate mutations that often result in increased transmissibility, pathogenicity, and avoidance of neutralizing antibody responses (Chu et al., 2020; Lin et al., 2021; Liu et al., 2021a; Liu et al., 2021b; McCallum et al., 2021; Mlcochova et al., 2021; Saito et al., 2021; Sheikh et al., 2021; Singh et al., 2021). For example, the Delta VOC (B.1.617.2), which is associated with a higher risk of hospitalization, intensive care unit (ICU) admission, and mortality (Lin et al., 2021; Sheikh et al., 2021) became the dominant strain worldwide within 6 months of its identification.

All VOCs to date have genetic changes in spike that are associated with in an increase in transmissibility, pathogenicity, and/or escape of antibody responses (reviewed in (Mengist et al., 2021; Park and Hwang, 2021; Thakur et al., 2021)). Spike, a trimeric type I fusion protein, targets cells for viral entry through a high-affinity interaction between the Receptor Binding Domain (RBD) and the cellular receptor Angiotensin Converting Enzyme (ACE) expressed on the target cell (Tortorici and Veesler, 2019). Spike was identified early in the pandemic as an attractive druggable target as knowledge of the related SARS-CoV-1 and MERS-CoV viruses could quickly be adopted to guide the development of SARS-CoV-2 prophylactic vaccines and recombinant antibody treatments (Salvatori et al., 2020). The current generation of vaccines and therapies have thus far proven to be extremely effective in limiting the incidence and severity of the disease (Boggiano et al., 2021; Liu et al., 2021d). However, current vaccines may not provide lifelong immunity against SARS-CoV-2. For example, the levels of protective antibodies and possibly cellular immunity resulting from mRNA based BNT162b2 vaccine has been shown to decline steadily (Kertes et al., 2021; Tartof et al., 2021) which is problematic since immunity against COVID-19 requires both humoral and cellular immunity (Altmann and Boyton, 2020; Karlsson et al., 2020; Sette and Crotty, 2021; Tan et al., 2021; Vabret et al., 2020). In addition, VOCs have accumulated a significant number of mutations, and those in spike are concerning since several vaccines use the sequence of the original Wuhan isolate. Indeed, a reduction in vaccine mediated-protection has been observed for Delta, Alpha, Beta, and Gamma to varying degrees (Hoffmann et al., 2021; Singh et al., 2021; Tegally et al., 2021; Thomson et al., 2021; Tregoning et al., 2021; Wang et al., 2021a; Wang et al., 2021b). Evasion of cellular immunity for these VOCs is thought to play a minor role since T cell epitopes previously identified (reviewed in (Grifoni et al., 2020)) are so far largely conserved and a negligible impact on T cell responses has been reported (Gallagher et al., 2021; Keeton et al., 2021; Redd et al., 2021b; Tarke et al., 2021b). However, the continual evolution of SARS-CoV-2 could result in a more significant impact on cellular immunity.

The latest VOC, Omicron (B.1.1.529), spread around the world approximately 3 times faster than Delta to become the dominant strain within two months of its emergence. The number of mutations encoded in Omicron largely exceeds the number found in other VOCs. The density of genetic changes in the Receptor Binding Motif (RBM) of Omicron spike, the critical region of ACE2 binding, is particularly concerning as they have the potential to provide a selective advantage as well as adversely impact the efficacy of vaccines developed against the RBM of the Wuhan isolate. Accordingly, early studies report an increased binding affinity of the Omicron RBD to human ACE2 (Cameroni et al., 2021) as well as a reduction of neutralizing activity of RBM-directed monoclonal antibodies and/or sera from individuals vaccinated with mRNA-1273, BNT162b2, AZD1222, Ad26.COV2.S, Sputnik V and BBIBP-CorV (Cameroni et al., 2021; Cele et al., 2021; Liu et al., 2021c). Interestingly, individuals who were vaccinated with BNT162b and had a previous infection had a higher SARS-CoV-2 neutralization capacity (Cele et al., 2021). This observation is consistent with T cell immunity conferred from a previous infection being a necessary component for effective immunity against Omicron which has been previously reported for other SARS-CoV-2 variants (Gallagher et al., 2021; Keeton et al., 2021; Redd et al., 2021b; Tarke et al., 2021b). Indeed, vaccinated individuals with previous exposure have a T cell repertoire capable of cross-recognizing Omicron and other variants (Keeton et al., 2021; Tarke et al., 2021a). Thus, the identification of T cell antigens remains crucial for our understanding of COVID-19 as well as informing the next generation of vaccines and treatments (Sauer and Harris, 2020).

To identify additional SARS-CoV-2 antigens we utilized ATLAS™ which has been previously shown to identify CD4^+^ and CD8^+^ T antigens in both infectious disease pathogens and mutated human genomes (Davies et al., 2015; Lam et al., 2021; Li et al., 2012; Long et al., 2014; Picard et al., 2015). The ATLAS assay utilizes autologous immune cells to identify T cell responses in an unbiased and comprehensive manner within the context of the cell’s ability to internally process and present antigens for CD4^+^ and CD8^+^ recognition (Nogueira et al., 2018). Additionally, direct cytokine secretion measurements coupled with our statistical methods has demonstrated that ATLAS can identify antigens missed by other methods (Lam et al., 2021).

The in-depth profiling of SARS-CoV-2 recognition presented here identifies underappreciated antigenic hot-spots which may inform the design of next-generation COVID-19 vaccines. In addition, ATLAS could be used as an Immunomonitoring tool during the development of these new vaccines. Finally, these studies might help predict the impact of mutations encoded in subsequent VOCs which will likely continue to emerge in the future.

## Materials and Methods

### Patient enrollment, sample collection, and processing

Peripheral blood mononuclear cells (PBMCs) were enriched and frozen from consented donors and classified into three cohorts: unexposed, severe, and mild. For the exposed cohorts, subjects were PCR confirmed donors and samples were collected 1 week to 9 months post-disease resolution. Severe and mild donors were recruited in roughly equal numbers and categorized as severe if they required hospitalization and mild if they did not. Unexposed donor PBMCs were collected before mid-2019. Donor demographic information is summarized in S1 Table. All samples were purchased from vendors who adhered to ethical practices for donor consent, compensation, and sample collection.

### Design of the SARS-CoV-2 Plasmid Library

To generate a SARS-CoV-2 plasmid library for use in the ATLAS assay (described below and Fig 1), a consensus sequence was derived by aligning 1676 sequences available on NCBI as of May 2020 to the SARS-CoV-2 reference sequence (NC_045512) using the Geneious Prime software (Biomatters). A panel of DNA fragments were designed to profile the entire SARS-CoV-2 proteome along with several control polypeptides (S2 Table). For SARS-CoV-2 *orfs* E, ORF6, ORF7a, ORF7b, ORF8, and ORF10, a single fragment encoding the entire gene was designed. For orf1ab, spike, orf3a, M, and N, multiple fragments overlapping by 20 amino acids were tiled across the length of each gene. To ensure all possible antigens were profiled for orf1ab, the fragments were designed to span the junctions of the 16 NSPs encoded in the immature orf1ab polypeptide. This approach ensures all possible antigens would be encoded, as an epitope spanning and NSP junction has been reported (Mateus et al., 2020). As a negative control antigen, 42 N-terminal amino acids of the gene blFP-Y3 from *Branchiostoma lanceolatum*, which encodes the fluorescent protein mNeon Green (abbreviated NG), was chosen. Multiple overlapping fragments of IE63 from Varicella-zoster virus and pp65 from Human cytomegalovirus were also designed for positive controls. The DNA fragments were codon optimized prior to synthesis and cloning (Twist Biosciences) in-frame with a red fluorescent protein (Fresno RFP; Atum Bio).

**Figure 1.**
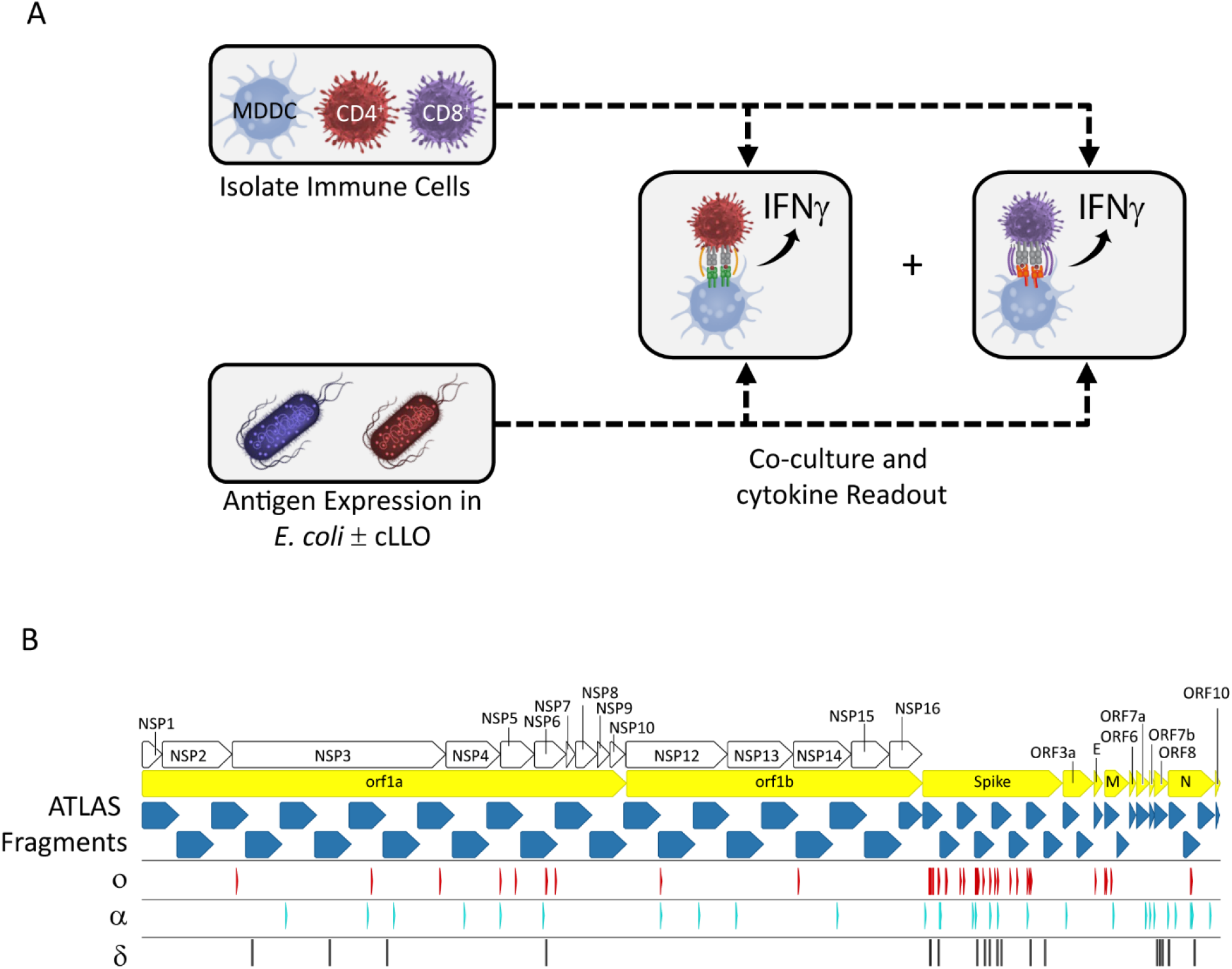
The ATLAS assay and library design. **(A)** Each putative antigen is expressed individually in two *E. coli* strains one with and one without cLLO (bottom left box), which are arrayed in expression libraries. Monocyte derived dendritic cells (MDDCs) as well as CD4^+^ and CD8^+^ T cells are isolated from peripheral blood and co-cultured overnight with the *E. coli* libraries and IFNγ output measured in each well. **(B)** Schematic representation of the library design depicting the location of the ATLAS Fragments (blue chevrons) in relation to the SARS-CoV-2 *orfs* (yellow chevrons) and orf1ab NSPs (white chevrons). Mutations from the Omicron (o), Delta (δ), and Alpha (α) VOCs are shown as red, cyan, and black barcodes, respectively.

### ATLAS Assay

The ATLAS assay was previously described (Lam et al., 2021). Briefly, the plasmid library was transformed by heat-shock into BL21(DE3) *E. coli* with and without a separate plasmid encoding a cytoplasmic variant of listeriolysin O (cLLO) for screening of CD8^+^ and CD4^+^ T cell antigens, respectively (Higgins et al., 1999). Subsequently, *E. coli* samples were cultured at 37°C in Luria Broth carbenicillin (100 μg/mL) ± Chloramphenicol (25 μg/mL) to select for transformed cells. Cultures were then diluted and cultured in selective media at 37°C for 3 hours and expression of polypeptides was induced with 3 mM IPTG and cultured over-night. Induced samples were fixed with formalin and then analyzed by flow cytometry to enumerate the bacteria and confirm antigen expression by fluorescence of the RFP reporter. Clones with confirmed expression of SARS-CoV-2 and positive control polypeptides were randomly re-arrayed in duplicate along with multiple replicates of the NG negative control into 384-well plates.

Magnetic microbeads (Miltenyi Biotec) were used to isolate CD14^+^ monocytes, CD8^+^ T cells, and CD4^+^ T cells from thawed PBMCs. The monocytes were cultured with IL-4 (13.8 ng/mL) and GM-CSF (20 ng/mL) for 6 days to generate monocyte-derived dendritic cells (MDDC). The CD8^+^ and CD4^+^ T cells were expanded with anti-CD3/anti-CD28 microbeads (Gibco) and IL-2 (178.5 IU/mL) for 6 days prior to magnetic bead removal and an overnight rest. MDDCs were co-cultured with the *E. coli* libraries for 2 hours to allow for processing and presentation of antigens. The plates were then washed prior to addition of CD8^+^ or CD4^+^ T cells to the appropriate *E. coli* library. After an overnight co-culture, IFNγ in the supernatant was measured using the MSD immunoassay according to the manufacturer’s instructions (Meso Scale Discovery).

### Data Analysis

Separate MSD readouts for CD4^+^ and CD8^+^ T-cells were generated from the ATLAS assay for each subject in the study. Quality checks were performed for each subject based on outlier detection, variability in dataset, plate controls, and completeness of dataset. The cytokine values within each plate were then normalized to NG using the formula below. Fragments with a mean normalized value > 1.75 were considered responses and if the response rate of all donors was between 50-74% or 75-100% it was either classified as common and frequent, respectively.

Normalization to NG:

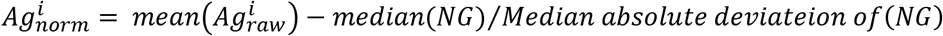

Where:

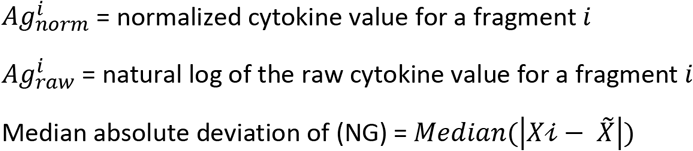

## Results

### ATLAS Identified Common and Frequent T Cell Antigens

Profiling of the exposed and unexposed cohorts using the ATLAS library depicted in Fig 1B revealed a wide breadth of responses throughout the SARS-CoV-2 proteome with every fragment producing a CD4^+^ and/or CD8^+^ response (Fig 2). Responses to ATLAS fragments were also observed for both T cell types in all *orfs* except for orf7b to which no donor had a CD8^+^ response. However, there was a skew towards S, M, and N as the number of responses per fragment for these *orfs* was larger (CD4^+^ = 11 and CD8^+^ = 11) than the rest of the proteome (CD4^+^ = 7 and CD8^+^ = 7). Biases were also observed when comparing the response profiles of the T cell subsets. For example, several *orfs* (orf3a, E, M, ORF7b, ORF8, and ORF10) and the C-Terminus of N were almost exclusively CD4^+^ targets. Additional biases were observed for orf1ab, CD4^+^ responses were localized to a few fragments in the orf1a region (NSPs 1-10) but were spread throughout the orf1b region (NSPs 12 – 16) while the reciprocal pattern was observed for CD8^+^ cells.

**Figure 2.**
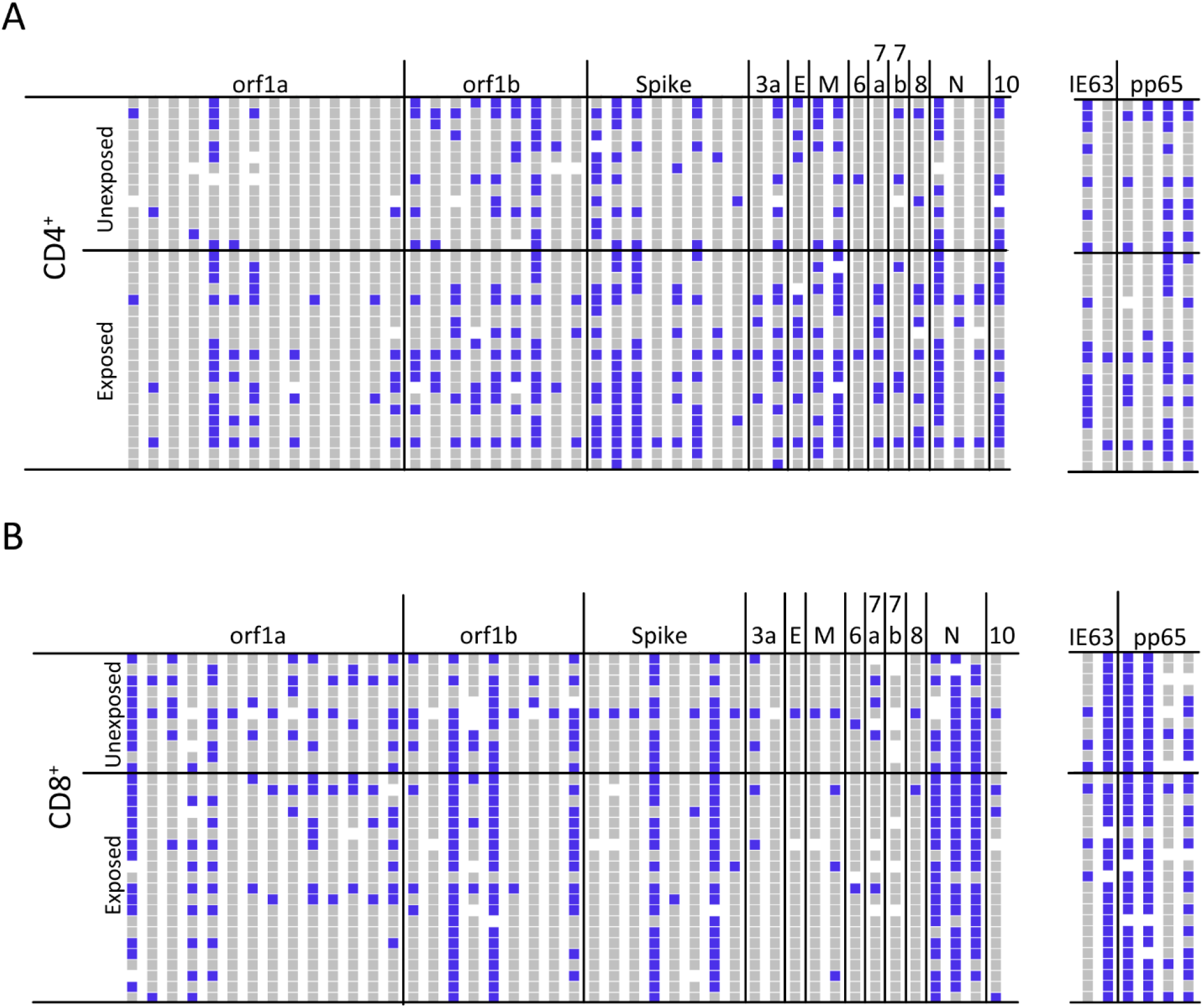
ATLAS identifies common and frequent SARS-CoV-2 antigens. **(A)** The CD4^+^ T cell response profiles to SARS-CoV-2 (left) and control antigens (right) for unexposed (N = 14) and exposed (N = 20) donors. Responses are blue, non-responses are grey, and white indicates not tested. Each column represents an ATLAS clone, and each row is a donor. Fragments and donors are grouped by *orf* and cohort, respectively. **(B)** The CD8^+^ response for unexposed (N = 11) and exposed (N = 21) donors are reported similarly.

Several common and frequent CD4^+^ and/or CD8^+^ SARS-CoV-2 antigens were identified in both cohorts (Fig 2 and S3 Table). Common and frequent antigens were identified in orf1ab, N, and spike whereas M and ORF10 only encoded common CD4^+^ antigens. Profiling of orf1ab identified a common CD4^+^ antigen in both orf1a and orf1b regions whereas frequent CD8^+^ antigens were identified in both regions along with a common antigen in orf1b.

### Unexposed Donors Often Responded to Multiple SARS-CoV-2 Antigens

When analyzing the response pattern of the unexposed cohort, multiple CD4^+^ (median = 6) and CD8^+^ (median = 8) responses were detected with all donors having at least one response (Fig 2). A wide breadth of responses was also observed in this cohort, with 72% and 90% of the fragments inducing at least one CD4^+^ and CD8^+^ response, respectively. Unexpectedly, a comparison of unexposed and exposed profiles revealed both cohorts responding to similar regions of the proteome at comparable frequencies (S3 Table). The strong similarities in response profiles resulted in the inability of our statistical analyses to stratify donors based on SARS-CoV-2 exposure or disease severity (data not shown). Although multiple studies observed the separation of exposed and unexposed cohorts, some unexposed donors in these studies responded to SARS-CoV-2 antigens potentially because of pre-exposure to related coronaviruses (Bacher et al., 2020; Le Bert et al., 2020; Sette and Crotty, 2020). As discussed in more detail below, ATLAS may be fine-tuned to identify otherwise infrequent cross-reactive antigens which might be missed by other methods, and which resulted in the absence of stratification in this study.

### The Spike ATLAS Response Profile Offers Insights into T cell Recognition of the RBM Domain

The spike peptide fragments were designed such that S_2-178 and S_159 encode the N terminal domain (NTD), S_316-492 and S_473-649 span the RBD/RBM, and the remaining 4 fragments were tiled across the C-terminus (Fig 3A). When mapping the position of the Alpha and Delta mutations onto these fragments, the polymorphisms are primarily located in the first half of the NTD and the central portion of the protein (*i.e*., the region encompassing the RBM and S1/S2 domains), however, neither variant had more than 3 mutations in any fragment (S3 Table). Conversely, the number of Omicron mutations mapped to each fragment outnumbered mutations from the other VOCs except for the last fragment which had none. In addition, 12 Omicron mutations mapped to fragments which encode the RBD (S_316-492 and S_473-649) with a particularly striking density in the second half of the RBM, the region critical for ACE2 binding. All eight of these mutated RBM residues are encoded in S_473-649 whereas 3 are in S_316-492 (Fig 3B).

**Figure 3.**
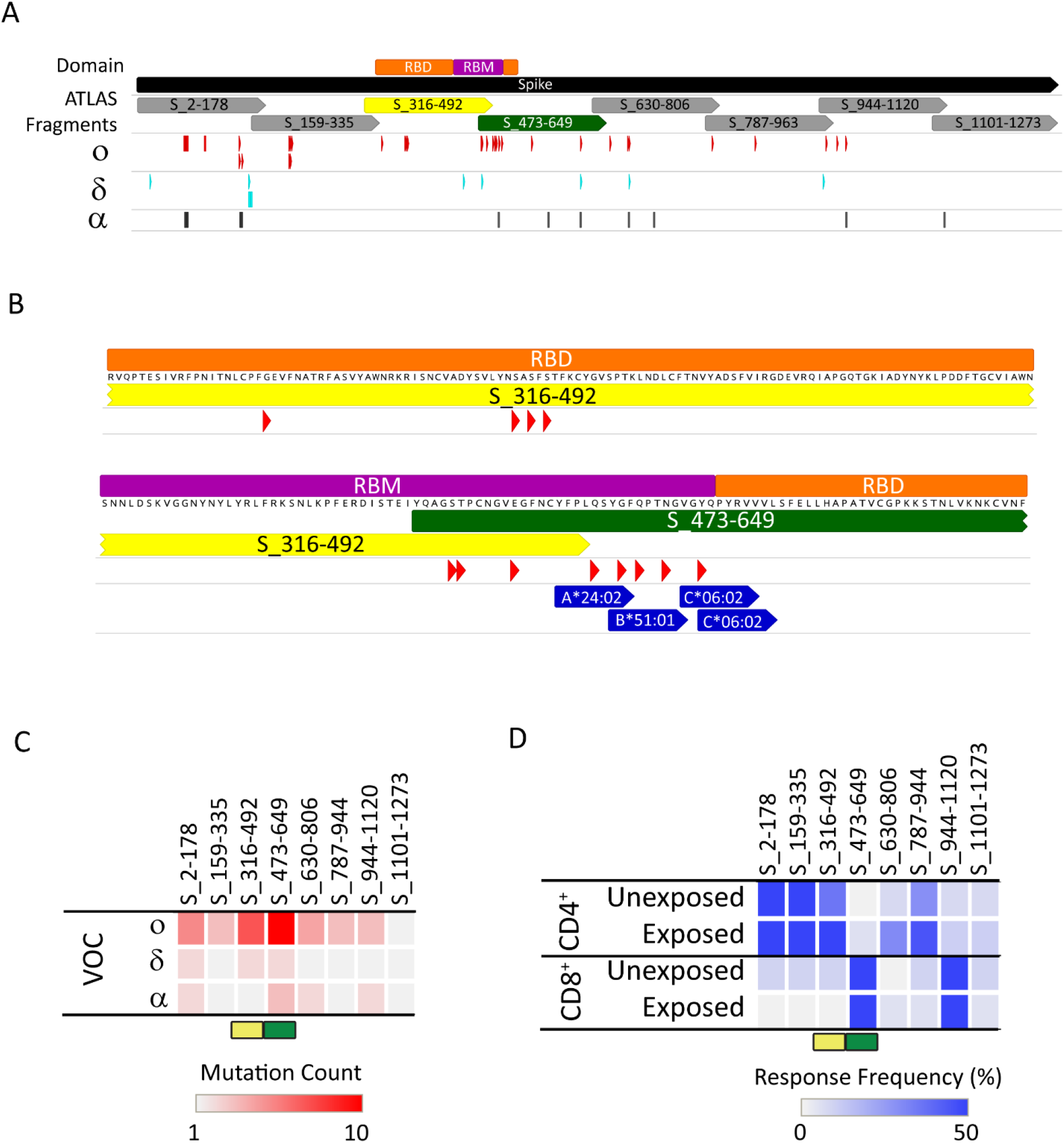
CD4^+^ and CD8^+^ T cells target complementary portions of the spike RBM. **(A)** Schematic representation of the ATLAS fragments (gray, yellow, and green chevrons), RBD (orange rectangle), and RBM (purple rectangle) in relation to the SARS-CoV-2 spike protein (black chevron). The position of the Omicron (ο), Delta (δ), and Alpha (α) VOC mutations are indicated by the red, cyan, and black barcodes, respectively. **(B)** Close up of the RBD domain region residues 319-437 and 438-541 are shown top and bottom, respectively. The RBD, RBM, and Omicron mutations are shown using the same color scheme. Also indicated, are previously reported CD8^+^ epitopes with known HLA restrictions encoded in the RBM region (blue chevrons). **(C and D)** Heatmaps indicating the number of mutations from α, δ, and ο VOCs encoded in each ATLAS fragment **(Panel C)** and the percentage of responding donors to each arranged by T cell subset and cohort **(Panel D)**. The rectangles highlight the fragments spanning the RBD and RBM domains S_316-492 (yellow) and S_473-649 (green).

As observed in the rest of genome, the ATLAS recognition profile of unexposed and exposed individuals was overlapping (Fig 3C). However, when comparing T cells subsets, a complementary pattern was observed. The CD8^+^ T cells from nearly all donors (94%; 30/32 donors) recognized at least one spike fragment with 75% recognizing at least two (24/32 donors). Analogously, 94% of donors (32/34) displayed at least 1 CD4^+^ response and 68% (11/34 donors) had at least 2. The majority of CD8^+^ responses were localized to fragments S_473-649 and S_944-1120 which were identified as frequent antigens and encode the RBM and the HR1/CH domains, respectively. Although, CD4^+^ T cell responses were found throughout the length of the protein, few class II responses were observed at the C-terminal end of the protein. Conversely, donors often responded to the N-terminus of spike with the first three fragments identified as common antigens.

## Conclusion

Most of the studies reported so far used *in silico* tools to predict potential antigenic peptides or used overlapping peptides. Using these approaches, approximately 1500 SARS-CoV-2 derived T cell epitopes have been described (reviewed in (Grifoni et al., 2021)). To further enhance our understanding of the cellular responses to SARS-CoV-2, we deployed ATLAS, which is an *ex vivo* assay capable of identifying CD4^+^ and CD8^+^ T cell antigens without prior knowledge of the donors’ HLA repertoire or putative antigenic sequences. ATLAS capitalizes on natural processing and presentation pathways (reviewed (Rock et al., 2016)) and can use peptides hundreds of amino acids long. As a result, it maximizes the ability of the immunoproteasome to generate and present optimally processed peptides compared to assays which rely exclusively on exogenous loading of synthetic peptides. Together, these factors could have contributed to the identification of additional ross-reactive antigens in the same regions which resulted in the homogenizing of response profiles and the absence of patterns which differentiated donors based on exposure or disease severity (refer to Fig 2).

ATLAS detected responses throughout the SARS-CoV-2 proteome along with several regions being characterized as common or frequent antigens. These antigenic regions corroborate and build upon the previously reported immunodominant regions derived through a meta-analysis of twenty-five SARS-CoV-2 antigen identification studies (Grifoni et al., 2021). The immunodominant CD8^+^ antigenic regions identified in that meta-analysis tended to be evenly distributed with a higher percentage of responding donors consistent with our observation that the most frequent ATLAS-identified antigens were class I restricted. Moreover, analysis of the ATLAS profiles revealed an enrichment of common and frequent antigens in structural protein *orfs*, but also identified several antigens in NSPs known to also be broadly recognized. Intriguingly, ORF10 was identified as a common CD4^+^ antigen in the unexposed cohort, but the source(s) of cross-reactive antigen(s) is unclear. Bioinformatic analysis did identify ORF10 homologs in bat and pangolin viruses as well as a truncated sequence in SARS-CoV-1 (Cagliani et al., 2020; Liu et al., 2020), but it seems unlikely that cross-reactive antigen(s) from these sources alone could account for the observation reported here. Since ORF10 has been implicated in suppressing the innate immune response (Li et al., 2021) further investigation into the origin of this common antigenic region is warranted. While we observed significant overlap in the dominant antigenic regions of SARS-CoV-2 antigens identified by ATLAS and previous studies, inconsistencies were still apparent. For example, ATLAS rarely detected a CD8^+^ response to the M protein, but the previous meta-analysis revealed several regions of immunodominance (Grifoni et al., 2021). Differences in the delivery of antigens (*i.e*., internal processing vs. exogenous loading) as well as sample sizes, *orfs* profiled, HLA alleles considered, and duration since symptom onset, *etc*. could all have significantly contributed to the differences observed in the T cell recognition landscape of SARS-CoV-2 across different studies.

The work presented here allowed us to make several useful observations that may guide understanding the impact that Omicron and possibly future VOCs mutations have on the ability of CD4^+^ and CD8^+^ T cells to recognize spike. The S_316-492 fragment contains 3 of Omicron’s 8 RBM mutations, and the reference strain did not elicit any significant CD8^+^ T cell reactivity. Hence, it may be safe to assume that, unless they introduce novel antigens, mutations in this area will bear no functional significance for CD8^+^ T cells. In contrast, the CD8^+^ T cell immunoprevalent S_473-649 fragment contains all the Omicron RBM mutations, with 5 being uniquely encoded within a span of 13 amino acids. In the region spanning those 5 mutations (refer to Fig 3b), 4 class I epitopes with known HLA restrictions (A*24:02, B*51:01, and C*06:02) have been identified (Grifoni et al., 2021). While we acknowledge that Omicron mutations might adversely affect those epitopes in certain populations, additional targets encoded in spike would reduce the overall impact on cellular immunity. Indeed, there are several other epitopes with A*24:02 (25), B*51:01 (5), and C*6:02 (1) restrictions (Grifoni et al., 2021) outside of the RBD region which would not be affected by Omicron mutations.

Similarly, the ability of CD4^+^ T cells to recognize Omicron and promote antibodies to the RBM should not be significantly reduced to the Omicron variant, since all of Omicron’s RBM mutations are contained in a fragment that was not identified as being immunoprevalent in the reference strain (S_473-649). Of the remaining 4 RBD mutations, 3 are clustered within 5 amino acids (S_316-492) which might impact only a handful of potential HLA combinations. It is also possible that mutations could produce sequences with enhanced peptide processing and presentation resulting in the generation of beneficial epitopes. Therefore, the overall impact to both CD8^+^ and CD4^+^ recognition of spike, and the ability to mount an immune response to the RBM/RBD domain might be minimal.

Taken together, the data presented here suggests that, at the population level, the natural cellular immunity to Omicron, especially as it relates to the ability to recognize spike and contribute to the generation of neutralizing antibodies against the RBM, should not be significantly different between Omicron and the reference strain. At the time of submission of this manuscript, several reports confirmed this hypothesis (Gao et al., 2022; GeurtsvanKessel et al., 2021; Keeton et al., 2021; Madelon et al., 2021; Redd et al., 2021a; Tarke et al., 2021a), further validating ATLAS as a potential tool to predictively assess the impact of as well as monitoring cellular immune responses to current and novel VOCs.

## Supporting information

S1 Table Donor Information

S2 Table ATLAS Fragment DNA and Protein Sequences

S3 Table Mutations Response Freq

## Acknowledgements

The authors are grateful for the donors who provided samples for our research study. They also thank members of Genocea’s ATLAS team including James Loizeaux, Madison Milaszewski, Oscar Cabrera, and Matt Rafferty.

## Supporting information

S1 Table. Donor Information

S2 Table ATLAS Fragment DNA and Protein Sequences

S3 Table Mutations, Percent Responses, and Antigen Classification of ATLAS Fragments

## Author Contributions

**J. Foti:** Conceptualization, resources, data sourcing, formal analysis, supervision, validation, investigation, visualization, methodology, writing-original draft and -editing. **K. Lema:** Experimentation, resources, data sourcing, visualization, formal analysis, writing-review and editing. **J. Strickland:** Experimentation, resources, writing-review and editing. **E. Tjon:** Resources, data sourcing, formal analysis, validation, methodology. **A. Li:** Conceptualization, resources, sample sourcing. **A. Rivera:** Experimentation, resources. **Crystal Cabral:** Experimentation, resources. **L. Cormier:** Resources, sample sourcing. **L. Dowal:** Resources, supervision, editing. **S. Rao:** Conceptualization, supervision, resources, sample sourcing. **V. Vemulapalli:** Resources, supervision, data sourcing, formal analysis, methodology, writing-review and editing. **J. Flechtner:** Conceptualization, supervision, resources, methodology, writing-review and editing.

