## Supplementary material for "The ATLAS™ screening assay reveals distinct CD4^+^ and CD8^+^ SARS-CoV-2 antigen response profiles which have implications to Omicron cellular immunity": S1 Table Donor Information

**Supplementary Table 1: Donor Information**

| Donor ID | Cohort | Age | Sex | Ethnicity | Required Hospitalization | Screened $CD4^+$ | Screened $CD8^+$ | Collection Time Post-Diagnosis (Days) |
|---|---|---|---|---|---|---|---|---|
| LP-001 | Unexposed | U* | U | U | NA | Y | Y | NA |
| LP-002 | Unexposed | 33 | M | Caucasian | NA | Y | N | NA |
| LP-004 | Unexposed | 25 | M | Caucasian | NA | Y | Y | NA |
| LP-005 | Unexposed | 39 | F | Unknown | NA | Y | Y | NA |
| LP-006 | Unexposed | 54 | F | Caucasian | NA | Y | N | NA |
| LP-009 | Unexposed | 45 | F | Caucasian | NA | Y | Y | NA |
| LP-010 | Unexposed | 43 | M | Hispanic | NA | Y | Y | NA |
| LP-011 | Unexposed | 64 | M | Caucasian | NA | Y | Y | NA |
| LP-012 | Unexposed | 58 | M | Hispanic | NA | Y | Y | NA |
| LP-013 | Unexposed | 48 | M | Caucasian | NA | Y | N | NA |
| LP-014 | Unexposed | 66 | M | Caucasian | NA | Y | Y | NA |
| LP-015 | Unexposed | 56 | M | African American | NA | Y | Y | NA |
| HC-002 | Unexposed | 38 | M | Hispanic | NA | Y | Y | NA |
| HC-003 | Unexposed | 58 | M | African American | NA | Y | Y | NA |
| C19-1 | Exposed | 38 | F | Caucasian | No | Y | Y | 75 |
| C19-2 | Exposed | 63 | M | Caucasian | No | Y | Y | 146 |
| C19-6 | Exposed | 29 | F | Caucasian | No | Y | Y | 117 |
| C19-13 | Exposed | 34 | F | Caucasian | No | Y | Y | 56 |
| C19-14 | Exposed | 27 | M | Caucasian | No | Y | Y | 66 |
| C19-15 | Exposed | 61 | M | Caucasian | No | Y | Y | 63 |
| C19-17 | Exposed | 23 | F | Caucasian | No | Y | Y | 31 |
| C19-18 | Exposed | 42 | F | Caucasian | No | Y | Y | 34 |
| C19-19 | Exposed | 25 | M | Caucasian | No | Y | Y | 48 |
| C19-20 | Exposed | 52 | M | Caucasian | No | Y | Y | 80 |
| C19-16 | Exposed | 38 | M | Caucasian | Yes | Y | Y | 80 |
| C19-21 | Exposed | 56 | F | Caucasian | Yes | Y | Y | 52 |
| C19-22 | Exposed | 40 | M | Caucasian | Yes | Y | Y | 68 |
| C19-23 | Exposed | 39 | M | Black | Yes | Y | Y | 83 |
| C19-24 | Exposed | 47 | M | Hispanic | Yes | Y | Y | 32 |
| C19-31 | Exposed | 68 | M | Hispanic | Yes | Y | Y | 113 |
| C19-32 | Exposed | 48 | M | Hispanic | Yes | Y | Y | 108 |
| C19-33 | Exposed | 65 | F | Hispanic | Yes | Y | Y | 95 |
| C19-34 | Exposed | 61 | F | Hispanic | Yes | Y | Y | 116 |
| C19-35 | Exposed | 30 | M | Hispanic | Yes | Y | Y | 77 |
| C19-53 | Exposed | 30 | F | Hispanic | Yes | N | Y | 115 |

*U = unknown information
