## Supplementary material for "The ATLAS™ screening assay reveals distinct CD4^+^ and CD8^+^ SARS-CoV-2 antigen response profiles which have implications to Omicron cellular immunity": S3 Table Mutations Response Freq

**Supplementary Table 3: Mutations, Percent Responses, and Antigen Classification of ATLAS Fragments**

| | Mutations | | | $CD4^{+}$ | | | | $CD8^{+}$ | | | |
|---|---|---|---|---|---|---|---|---|---|---|---|
| | | | | % Responses | | | Common/ Frequent* | % Responses | | | Common/ Frequent* |
| Fragment | ο | δ | α | Unexposed | Exposed | Both | | Unexposed | Exposed | Both | |
| orf1ab_2-334 | | | | 8 | 5 | 6 | | 82 | 74 | 77 | F |
| orf1ab_315-647 | | | | 7 | 10 | 9 | | 18 | 5 | 9 | |
| orf1ab_628-960 | 1 | | | 0 | 0 | 0 | | 45 | 5 | 19 | |
| orf1ab_941-1273 | | | 1 | 8 | 0 | 3 | | 11 | 40 | 31 | |
| orf1ab_1254-1586 | | 1 | | 46 | 65 | 58 | C | 55 | 38 | 44 | |
| orf1ab_1567-1899 | | | 1 | 7 | 25 | 18 | | 9 | 0 | 3 | |
| orf1ab_1880-2212 | 1 | 1 | | 8 | 45 | 31 | | 20 | 10 | 13 | |
| orf1ab_2193-2525 | | 1 | 1 | 0 | 0 | 0 | | 9 | 10 | 9 | |
| orf1ab_2506-2838 | 1 | | | 0 | 16 | 9 | | 40 | 10 | 19 | |
| orf1ab_2819-3151 | | 1 | | 0 | 5 | 3 | | 27 | 33 | 31 | |
| orf1ab_3132-3464 | 2 | 1 | | 0 | 0 | 0 | | 18 | 10 | 13 | |
| orf1ab_3445-3777 | 2 | 1 | 1 | 0 | 0 | 0 | | 27 | 15 | 19 | |
| orf1ab_3758-4090 | | | | 0 | 10 | 6 | | 9 | 14 | 13 | |
| orf1ab_4071-4405 | | | | 8 | 26 | 19 | | 55 | 35 | 42 | |
| orf1ab_4382-4714 | | | | 36 | 32 | 33 | | 36 | 14 | 22 | |
| orf1ab_4695-5027 | 1 | 1 | | 21 | 15 | 18 | | 0 | 0 | 0 | |
| orf1ab_5008-5340 | | 1 | | 15 | 35 | 27 | | 73 | 95 | 88 | F |
| orf1ab_5321-5653 | | | | 14 | 26 | 21 | | 22 | 16 | 18 | |
| orf1ab_5634-5966 | | | | 36 | 40 | 38 | | 82 | 90 | 88 | F |
| orf1ab_5947-6279 | 1 | | | 38 | 32 | 34 | | 9 | 5 | 6 | |
| orf1ab_6260-6592 | | 1 | | 71 | 60 | 65 | C | 20 | 0 | 6 | |
| orf1ab_6573-6905 | | | | 8 | 5 | 6 | | 9 | 0 | 3 | |
| orf1ab_6886-7096 | | | | 0 | 15 | 9 | | 70 | 57 | 61 | C |
| S_2-178 | 5 | 3 | 2 | 62 | 55 | 58 | C | 9 | 0 | 3 | |
| S_159-335 | 3 | | | 50 | 75 | 65 | C | 9 | 0 | 3 | |
| S_316-492 | 7 | 2 | | 29 | 70 | 53 | C | 9 | 0 | 3 | |
| S_473-649 | 10 | 2 | 3 | 0 | 5 | 3 | | 82 | 81 | 81 | F |
| S_630-806 | 4 | 1 | 2 | 7 | 30 | 21 | | 0 | 5 | 3 | |
| S_787-963 | 3 | 1 | | 29 | 45 | 38 | | 9 | 5 | 6 | |
| S_944-1120 | 3 | 1 | 2 | 7 | 10 | 9 | | 82 | 90 | 87 | F |
| S_1101-1273 | | | 1 | 7 | 10 | 9 | | 9 | 5 | 6 | |
| Orf3a_2-148 | | 1 | | 0 | 20 | 12 | | 27 | 10 | 16 | |
| Orf3a_129-275 | | | | 38 | 45 | 42 | | 0 | 0 | 0 | |
| E | 1 | | | 21 | 32 | 27 | | 9 | 0 | 3 | |
| M_2-134 | 3 | 1 | | 36 | 45 | 41 | | 9 | 0 | 3 | |
| M_115-222 | | | | 43 | 68 | 58 | C | 9 | 14 | 13 | |
| ORF6 | | | | 7 | 5 | 6 | | 9 | 5 | 6 | |
| ORF7a | | 2 | | 0 | 40 | 24 | | 30 | 6 | 14 | |
| ORF7b | | 1 | | 15 | 20 | 18 | | 0 | 0 | 0 | |
| ORF8 | | 1 | 3 | 14 | 37 | 27 | | 9 | 5 | 6 | |
| N_2-154 | | 1 | | 62 | 70 | 67 | C | 56 | 86 | 77 | F |
| N_135-287 | 2 | 2 | 1 | 0 | 15 | 9 | | 100 | 86 | 90 | F |
| N_268-419 | | 1 | | 0 | 21 | 12 | | 82 | 81 | 81 | F |
| ORF10 | | | | 54 | 55 | 55 | C | 9 | 14 | 13 | |

Notes:

C (Common Antigen; 50 -74% of donors)

F (Frequent Antigen; 75 – 100% of donors)
